# Distinct glycerophospholipids potentiate Gsα-activated adenylyl cyclase activity

**DOI:** 10.1101/2022.02.23.481604

**Authors:** Anubha Seth, Marius Landau, Andrej Shevchenko, Sofia Traikov, Anita Schultz, Sherif Elsabbagh, Joachim E. Schultz

**Author notes:** both contributed equally (Anubha Seth, Marius Landau). Dept. of Pharmacology, Yale University School of Medicine, 333 Cedar Street, New Haven, CT 06520-8066, USA. Corresponding author: Dr. Joachim E Schultz, Pharmazeutisches Institut der Universität Tübingen, Auf der Morgenstelle 8, 72076 Tübingen, Germany.

## Abstract

Nine mammalian adenylyl cyclases (AC) are pseudoheterodimers with two hexahelical membrane domains which are isoform-specifically conserved. Previously we proposed that these membrane domains are orphan receptors (10.7554/eLife.13098; 10.1016/j.cellsig.2020.109538). Lipids extracted from fetal bovine serum at pH 1 inhibited several mAC activities. Guided by a lipidomic analysis we tested glycerophospholipids as potential ligands. Contrary to expectations we surprisingly discovered that 1-stearoyl-2-docosahexaenoyl-phosphatidic acid (SDPA) potentiated Gsα-activated activity of human AC isoform 3 seven-fold. The specificity of fatty acyl esters at glycerol positions 1 and 2 was rather stringent. 1-Stearoyl-2-docosahexaenoyl-phosphatidylserine and 1-stearoyl-2-docosahexaenoyl-phosphatidylethanolamine significantly potentiated several Gsα-activated mAC isoforms to different extents. SDPA appears not interact with forskolin activation of AC isoform 3. SDPA enhanced Gsα-activated AC activities in membranes from mouse brain cortex. The action of SDPA was reversible. Unexpectedly, SDPA did not affect cAMP generation in HEK293 cells stimulated by isoproterenol, PGE_2_ and adenosine, virtually excluding a role as an extracellular ligand and, instead, suggesting an intracellular role. In summary, we discovered a new dimension of intracellular AC regulation by chemically defined glycerophospholipids.

## Introduction

cAMP is a universal regulator of numerous cellular processes (Dessauer, Watts et al., 2017, Schultz & Natarajan, 2013, Sinha & Sprang, 2006, Sunahara & Taussig, 2002). Its biosynthesis is via adenylyl cyclases. This report deals with the nine mammalian, membrane-bound pseudoheterodimeric ACs (mACs; reviewed in (Bassler, Schultz et al., 2018, Dessauer et al., 2017, Ostrom, LaVigne et al., 2022, Schultz & Natarajan, 2013). Currently, a direct regulation of mACs does not exist. The accepted regulation is indirect and includes *(i)* the extracellular activation of G-protein-coupled receptors, *(ii)* intracellular release of the Gsα subunit from a trimeric G-protein and, *(iii)*, as a last step mAC activation by the free α-subunit (Dessauer et al., 2017, Sadana & Dessauer, 2009). Secondarily, calmodulin, Ca^2+^ ions, G_βγ_ and phosphorylation are cytosolic effectors. In contrast, we recently have assigned a direct regulatory role mediated by the membrane domains of mACs acting as receptors (Beltz, Bassler et al., 2016, Seth, Finkbeiner et al., 2020). In this proposal the mAC receptors are comprised of the two hexahelical domains each connected to a cytosolic catalytic domain, C1 and C2, via highly conserved cyclase-transducing-elements (Dessauer et al., 2017, Seth et al., 2020, Ziegler, Bassler et al., 2017). The proposal for a receptor function is based on *(i)* the evolutionary conservation of the membrane anchors in an isoform-specific manner for more than 0.5 billion years (Bassler et al., 2018), *(ii)* on highly conserved cyclase-transducing-elements (Ziegler et al., 2017), and *(iii)* on catalytic domains conserved from cyanobacteria to mammals (Bassler et al., 2018, Kanacher, Schultz et al., 2002, Linder & Schultz, 2003, Seth et al., 2020). Recently we reported ligand-mediated inhibition of a Gsα-activated mAC2 in a chimera in which the AC membrane domains were replaced by the hexahelical quorum-sensing receptor CqsS from *Vibrio sp.* which has a known lipophilic ligand, cholera-auto-inducer-1 (Beltz et al., 2016, Ng, Wei et al., 2010, Seth et al., 2020).

In an initial approach to identify ligands for the mAC receptors we used fetal bovine serum (FBS) which had been shown to contain inhibitory components (Seth et al., 2020). Eliminating peptides or proteins as possible ligands we fractionated lipids by extraction with chloroform/methanol at different pH values (Bligh & Dyer, 1959). Expecting to isolate inhibitory components we report the most surprising discovery that 1-stearoyl-2-docosahexaenoyl-phosphatidic acid (SDPA) potentiated Gsα-activated mAC3 activity up to 7-fold. The actions of SDPA resemble, to a limited extent, those of the plant diterpene forskolin (Dessauer et al., 2017). The data establish a new layer of direct mAC regulation and emphasize the importance of glycerophospholipids (GPLs) in regulation of intracellular cAMP generation.

## Results

### Lipids as possible mAC effectors

In exploratory experiments, we extracted lipids from FBS with chloroform/methanol at pH 1, pH 6, and pH 14 (Bligh & Dyer, 1959). After evaporation of solvent the solid residues were dissolved in DMSO and tested against human mAC isoforms 1, 2, 3, 5, 6, 7, 8, and 9 in membrane preparations from HEK293 cells transfected with the respective ACs (Table 1). The pH 1 extract inhibited Gsα-activated AC 1, 2, 5, 6, and 7 to different extents. The pH 6 and pH 14 extracts appeared to enhance Gsα-activated AC isoforms 2, 3, 8, and 9 (table 1).

**Table 1.** Lipid Extraction of FBS with Chloroform / Methanol.

**% Gsα activated Adenylyl Cyclase Activity**
|  | pH 14 | pH 6 | pH 1 |
| --- | --- | --- | --- |
| **hAC1** | **99±5** | **91±5** | **25±1** |
| **hAC2** | **176±8** | **158±4** | **149±9** |
| **hAC3** | **183±40** | **175±41** | **31±11** |
| **hAC5** | **98±7** | **123±12** | **50±7** |
| **hAC6** | **91±4** | **89±2** | **57±4** |
| **hAC7** | **140±6** | **131±4** | **79±6** |
| **hAC8** | **192±11** | **193±7** | **115±4** |
| **hAC9** | **201±22** | **251±12** | **140±12** |

2mL FBS were extracted with chloroform/methanol (1:2) according to (Bligh & Dyer, 1959). The organic phase was evaporated and the residue was dissolved in 35 μl DMSO. Adenylyl cyclases were activated by 600 nM Gsα and 33 nl of the DMSO extracts were added. Basal AC activities were in the order of the table: 0.11, 0.43, 0.02, 0.4, 0.16, 0.02, 0.33, and 0.04 nmol cAMP·mg^-1^·min^-1^, respectively. 600 nM Gsα-activated activities were 0.49, 1.31, 0.36, 2.23, 0.71, 0.25, 3.16 and 1.67 nmol cAMP·mg^-1^·min^-1^. n = 4 to 12.

We then carried out a lipidomic analysis with the pH 1 and the pH 6 fractions (Matyash, Liebisch et al., 2008, Vvedenskaya, Rose et al., 2021). Based on previous data we expected potential ligands which inhibit Gsα-activated mAC activities and concentrated on lipids present in the pH 1 fraction (Seth et al., 2020). Apart from several minor constituents from different lipid classes the major constituents in the acidic fraction were phosphatidic acids, phosphatidylcholine, phosphatidylethanolamine and phosphatidylserines (see Appendix Fig. 1 and 2). Next we examined the effect of commercially available bulk lipids on Gsα-activated mACs. Egg phosphatidic acids significantly stimulated, whereas other bulk lipids such as egg and liver phosphatidylcholine, brain gangliosides, sulfatides and cerebrosides had no significant effects. The lipidomic analysis showed that highly unsaturated fatty acids such as arachidonic acid and docosahexaenoic acid are prominent acyl substituents in phosphatidic acids (Appendix Fig. 2). These acyl residues are only minor components in the tested egg or liver phosphatidic acids. Therefore, we assayed commercially available synthetic GPLs containing polyunsaturated fatty acids as acyl substituents. The general structure of glycerophospholipids is shown below (see Appendix Table 1 for a complete list of lipids examined in this study).

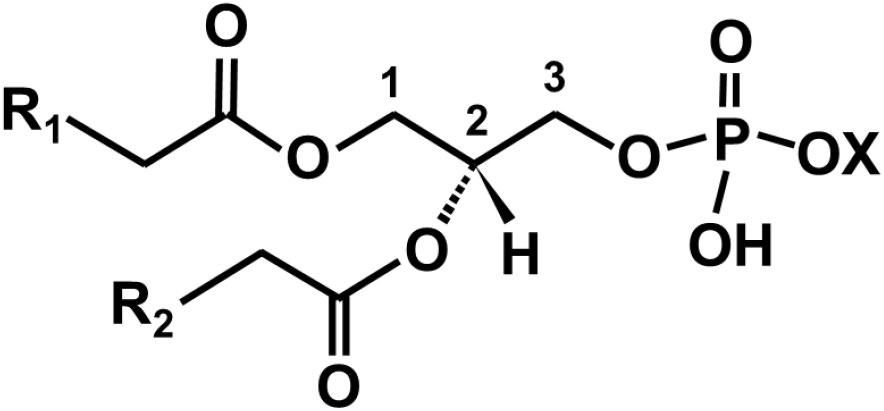
Basic structure of glycerophospholipids: R_1_and R_2_ are fatty acyl residues esterified at glycerol positions 1 and 2; X can be a proton H^+^ as in phosphatidic acid, choline (phosphatidylcholine), serine (phosphatidylserine), glycerol (phosphatidylglycerol), or ethanolamine (phosphatidylethanolamine).

The assays used membranes containing human mAC isoforms expressed in HEK293 cells. The mACs were activated by 600 nM of a constitutively active Gsα (Q227L, here termed Gsα) because we expected to characterize an inhibitory input (Graziano, Freissmuth et al., 1991, Seth et al., 2020). Most surprisingly, we discovered that 1-stearoyl-2-docosahexaenoyl-phosphatidic acid (SDPA) potentiated mAC3 up to 7-fold above the 16-fold activation already exerted by 600 nM Gsα alone (Fig. 1). The EC_50_ of SDPA was 0.9 μM. In the absence of Gsα 10 μM SDPA had no significant effect (Fig. 1). As far as the synergism between Gsα-activated mAC3 is concerned the effect of SDPA was reminiscent of the known cooperativity between forskolin and Gsα activated mACs (Dessauer et al., 2017).

**Figure 1.**
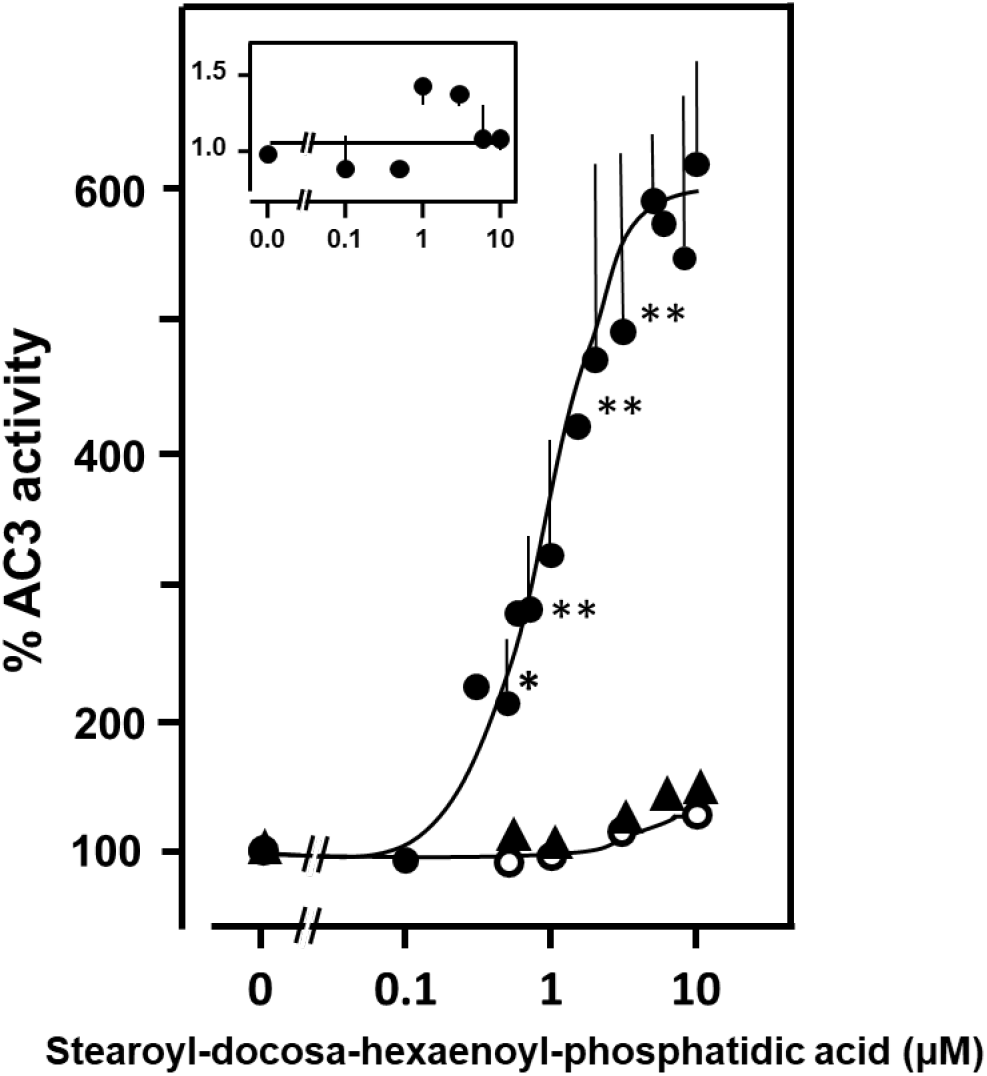
1-Stearoyl-2-docosahexaenoyl-phosphatidic acid concentration-dependently potentiates mAC3 activated by 600 nM Gsα (filled circles). 100 % Gsα-activated mAC3 activity corresponded to 707 ± 187 pmoles cAMP•mg^-1^•min^-1^. Basal mAC3 activity is not significantly affected by SDPA (open circles; 100 % basal activity corresponds to 34 pmoles cAMP•mg^-1^•min^-1^). Triangles: Effect of SDPA on the C1-C2 soluble AC construct activated by 600 nM Gsα (Basal activity was 12 pmol cAMP•mg^-1^∂min^-1^. 600 nM Gsα activated activity was 150 pmol cAMP•mg^-1^•min^-1^ corresponding to 100 %). Insert: Activity of the mycobacterial AC Rv1625c is unaffected by SDPA (activity was 23 nmoles cAMP•mg^-1^•min^-1^). Data were normalized to respective 100 % activities. Significances in a two-tailed t-test: *: p<0.05; **: p<0.01 compared to 100% activity. For clarity, not all significances are marked. N = 4-6; error bars denote S.E.M.’s.

Does the action of SDPA require a membrane-anchored AC holoenzyme or is the activity of a Gsα-activated C1/C2 catalytic dimer potentiated as well? We produced a soluble active AC construct connecting the catalytic C1 domain of mAC1 and the C2 domain of mAC2 by a flexible linker (Tang & Gilman, 1995). The construct was expressed in *E. coli* and purified via its His_6_-tag. It was activated 12-fold by Gsα (from 12 to 150 pmol cAMP•mg^-1^•min^-1^). SDPA up to 10 μM did not affect basal activity and failed to significantly enhance Gsα-activated activity of the chimera. We tentatively conclude that the SDPA action requires membrane anchoring of mACs.

We investigated whether SDPA affects the activity of a Gsα-insensitive membrane-bound bacterial AC. We used the mycobacterial AC Rv1625c, a monomeric progenitor of mammalian mACs, which has a hexahelical membrane domain and is active as a dimer (Guo, Seebacher et al., 2001). The activity of the Rv1625c holoenzyme was unaffected by SDPA (Fig. 1 insert). The particular intrinsic properties of the mammalian membrane domains in conjunction with Gsα-activation may be required to confer SDPA sensitivity.

Next, we examined which kinetic parameters are affected by SDPA. For mAC3, the enzymatic reaction rates ± SDPA were linear with respect to protein concentration and time up to 30 min. The Km for substrate ATP (0.1 mM) was unaffected. The most striking effect of SDPA was the increase in Vmax (from 4 to 8 nmol cAMP•mg^-1^•min^-1^). Concentration-response curves for Gsα in the presence of different SDPA concentrations showed that the affinity of mAC3 for Gsα was significantly increased (Fig. 2). Most likely, Gsα and SDPA act at distinct sites of the protein and potentiation by SDPA is due to concerted structural interactions, reminiscent of the cooperativity between Gsα and forskolin (Dessauer et al., 2017).

**Figure 2.**
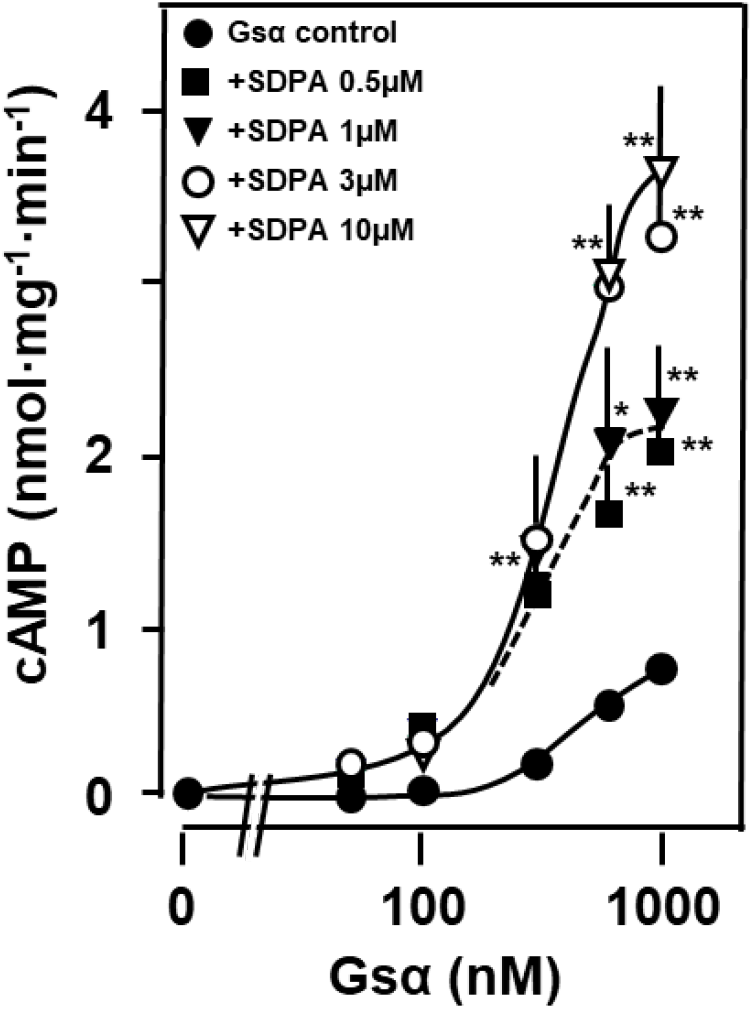
SDPA increases the affinity of mAC3 for Gsα. The EC_50_ concentration for Gsα in the absence of SDPA was 518 nM and in the presence it was 336 ± 29 nM (p<0.02; n=5-6). Basal AC3 activity was 27 ± 21 pmoles cAMP·mg^-1^•min^-1^; 1000 nM Gsα increased mAC3 activity to 791± 128 pmoles cAMP•mg^-1^·min^-1^). Significances: *: p<0.05; **: p<0.01 compared to corresponding activities without SDPA. n = 5-6; error bars denote S.E.M. Often, error bars did not exceed the symbol size.

### Specificity of 1- and 2-acyl substituents in phosphatidic acid

Phosphatidic acid is the simplest GPL consisting of a glycerol backbone to which two fatty acids and phosphoric acid are esterified. At physiological pH it carries about 1.5 negative charges. Generally, at positions 1 and 2 of glycerol a variety of fatty acyl residues haves been identified. We examined the biochemical specificity of the fatty acyl substituents in phosphatidic acid (Fig. 3).

**Figure 3.**
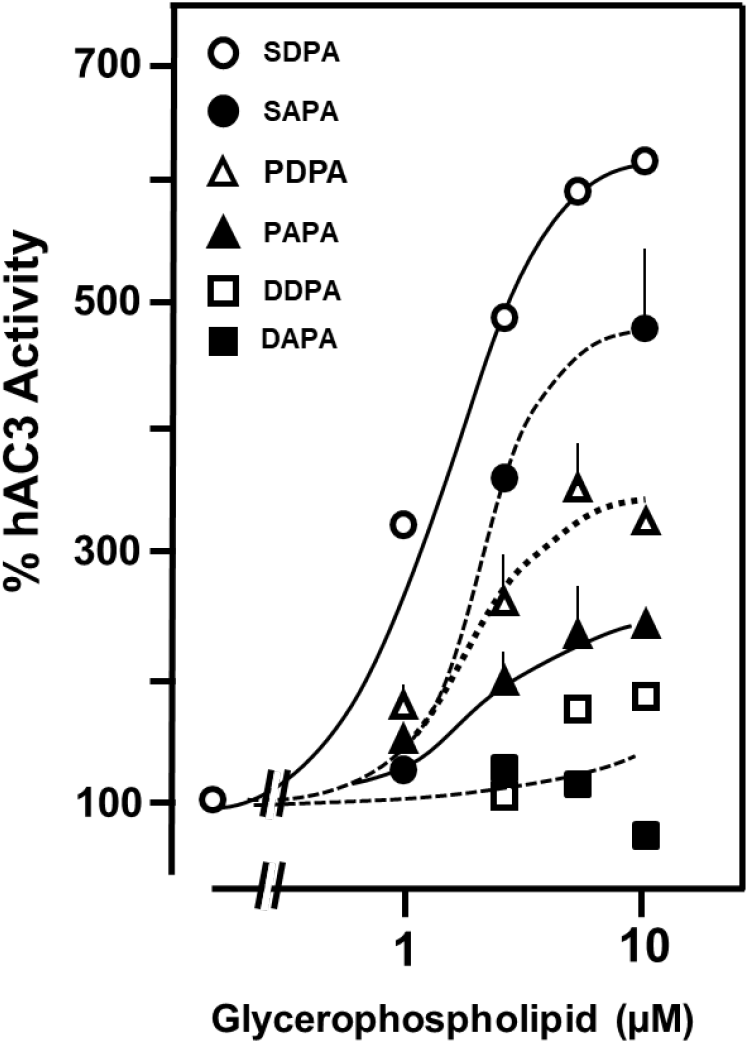
Specificity of fatty acyl esters in phosphatidic acids for potentiation of Gsα-activated mAC3. 600 nM Gsα-activated activity (100%) was 446 pmol cAMP•mg^-1^•min^-1^ (basal mAC3 activity was 15.2 pmol cAMP•mg^-1^•min^-1^). Abbreviations: SAPA, 1-stearoyl-2-arachidonoyl-phosphatidic acid; PDPA, 1-palmitoyl-2-docosahexaenoyl-phosphatidic acid; PAPA, 1-palmitoyl-2-arachidonoyl-phosphatidic acid; DDPA, di-docosahexaenoyl-phosphatidic acid; DAPA, di-arachidonoyl-phosphatidic acid. The EC_50_ concentrations were 4.8, 1.3, and 1.4 μM, for SAPA, PDPA, and DDPA, respectively (differences not significant; n=3). Error bars denote S.E.M. For comparison, a curve presenting SDPA is included.

10 μM 1-Stearoyl-2-arachidonoyl-phosphatidic acid (SAPA) potentiated Gsα-activated mAC3 about 5-fold (EC_50_ = 4.8 μM; Fig. 3). Exchanging the stearic acid at position 1 by a palmitic acid, i.e. 1-palmitoyl-2-docosahexaenoyl-phosphatidic acid (PDPA) reduced activity by about 50 % compared to SDPA (EC_50_ = 1.3 μM). Strikingly, the corresponding 1-palmitoyl-2-arachidonoyl-phosphatidic acid (PAPA) lost about 70% of activity compared to SDPA (Fig. 3), highlighting the structural contribution of the 1-fatty acyl substituent to biochemical activity. The importance of the substituent at position 1 was further emphasized when assaying 1, 2-di-docosahexaenoyl-phosphatidic acid (DDPA). The efficiency was reduced by 80 % compared to SDPA (Fig. 3). The EC_50_ for DDPA was 1.4 μM. Even more drastic was the absence of an effect using 1, 2-di-arachidonoyl-phosphatidic acid (DAPA; Fig. 3). Expectedly then, 1-stearoyl-2-linoleoyl-phosphatidic acid was inactive (not shown). The data show a remarkable positional specificity for the 1- and 2-acyl substituents of the glycerol backbone and indicate a specific and concerted interaction between the fatty acyl esters. The specificity of fatty acyl-substitution also strongly indicated that SDPA is not acting in its property as a general membrane GPL because other phosphatidic acids should be equally suitable as membrane lipids. Further, the peculiar biochemical properties of SDPA in its relation with AC isoforms suggest that the negative charges of phosphatidic acid are not sufficient to determine specificity, but that the lipid substitutions on position 1-as well as 2-probably are equally important.

### Head group specificity of glycerophospholipids

The next question is whether 1-stearoyl-2-docosahexaenoyl-GPLs with different head groups might affect Gsα-activated mAC3. First, we replaced the phosphate head group in SDPA by phosphoserine generating SDPS. This greatly reduced potentiation of Gsα-activated mAC3 activity (2.8-fold potentiation; Fig. 4). A concentration-response curve of SDPS with mAC3 showed that the EC_50_ concentration was similar to that of SDPA (1.2 vs 0.9 μM; n=6-9; n.s.), but its efficacy is significantly lower suggesting that identical binding sites are involved.

**Figure 4.**
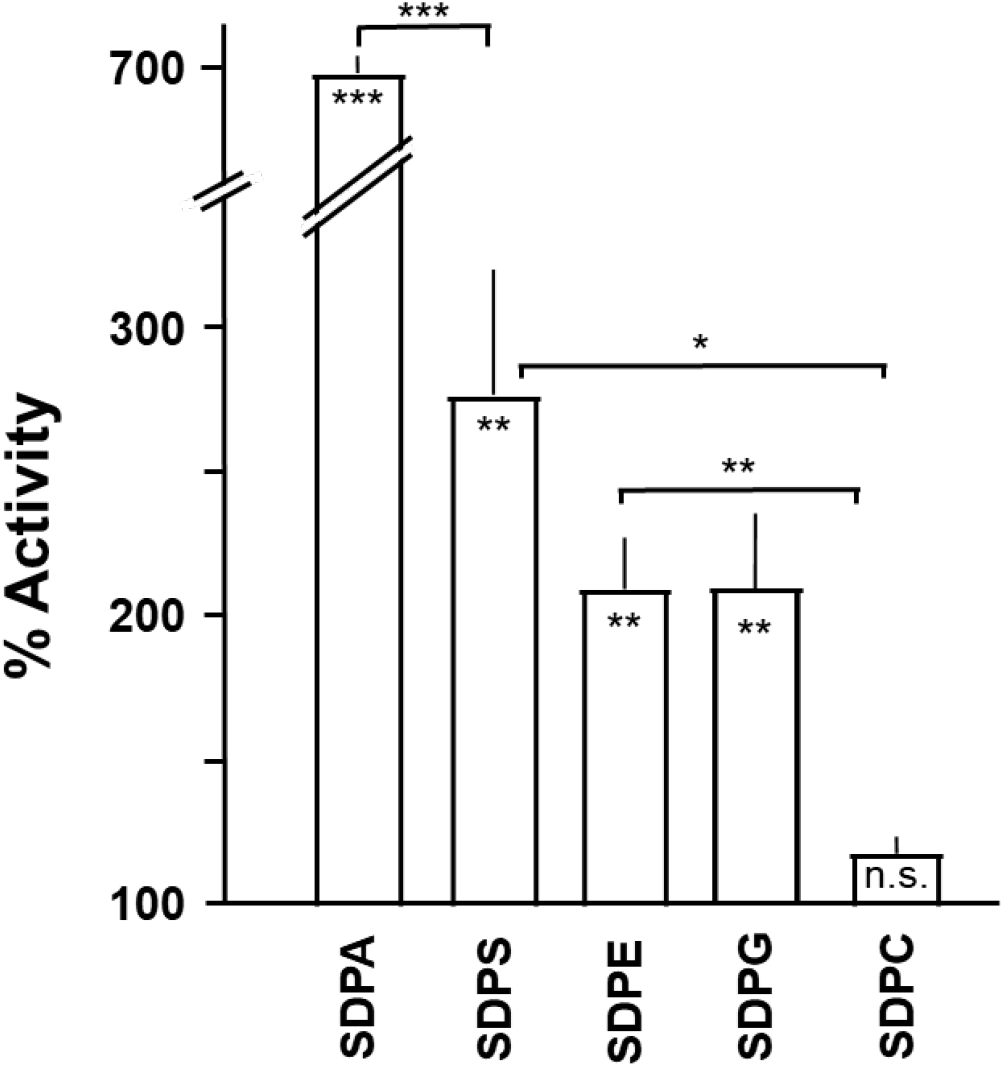
Head group specificity of glycerophospholipids enhancing Gsα-activated mAC3 activity. Basal mAC3 activity was 0.03. Gsα-stimulated activity was 0.56 nmol cAMP•mg^-1^•min^-1^ (corresponding to 100 %). Concentration of lipids was 10 μM. Error bars denote S.E.M. Significances: *: p< 0.05; **: p< 0.01; ***: p<0.001; n = 3-9.

We further used 1-stearoyl-2-docosahexaenoyl-ethanolamine (SDPE), 1-stearoyl-2-docosahexaenoyl-phosphatidylglycerol (SDPG) and 1-stearoyl-2-docosahexaenoyl-phosphatidylcholine (SDPC; Fig. 4). In this order, efficacy to enhance the Gsα-activated mAC3 declined, with SDPC having no significant effect (Fig. 4). The surprising specificity of the 1- and 2-fatty acyl-substituents of the glycerol backbone was emphasized once again when we used 1-stearoyl-2-arachidonoyl-phosphatidyl-ethanolamine (SAPE) and 1-stearoyl-2-arachidonoyl-phosphatidylcholine. In both instances biochemical activity was lost (not shown). Consequently, we did not further probe GPLs with differing fatty acyl combinations at the glycerol 1- and 2-positions because, as demonstrated, changes in acyl substitutions resulted in considerable reduction or loss of biological activity (see Fig 3).

### Effect of glycerophospholipids on Gsα-activated adenylyl cyclase isoforms

So far, we examined only the mAC3 isoform which showed a particularly high synergism between Gsα and SDPA. Does SDPA equally potentiate the Gsα-activated activities of the other mAC isoforms? More generally, do GPLs display an mAC isoform specificity in the regulation of intracellular cAMP biosynthesis? We expressed the nine human mAC isoforms in HEK293 and, first, tested how SDPA affected the Gsα-activated activities (Fig. 5).

**Figure 5.**
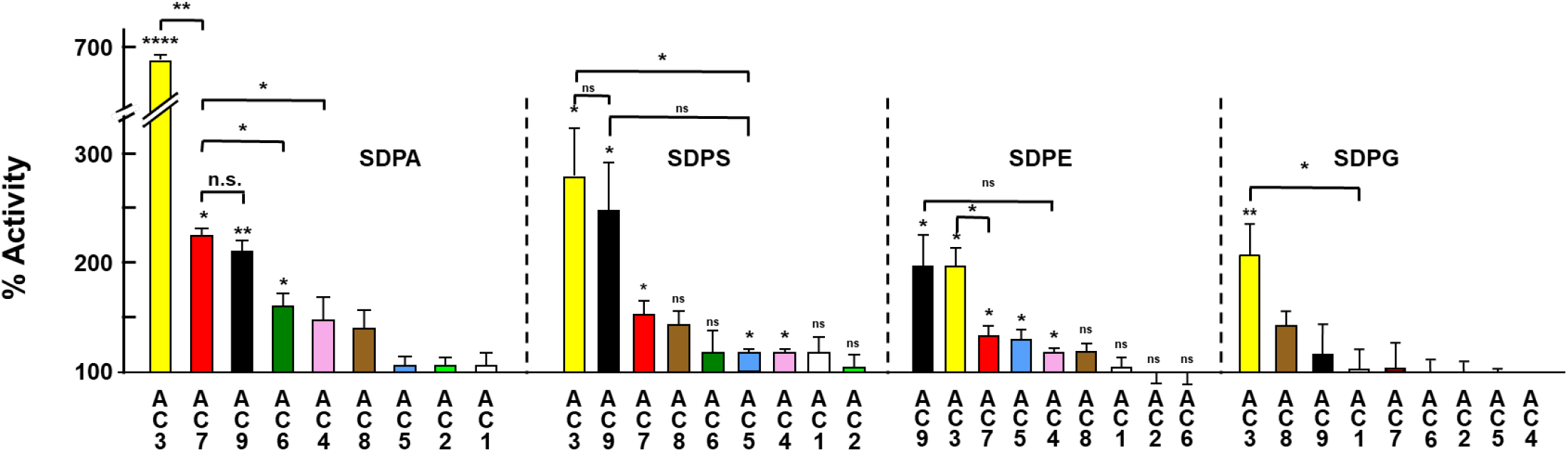
Effect of 10 μM of glycerophospholipids on various mAC isoforms activated by 600 nM Gsα. Basal activities and Gsα-activated activities are listed in Appendix Table 2). Error bars denote S.E.M. Significances: *: p< 0.05; **: p< 0.01; ***: p<0.001; ****: p<0.0001 compared to Gsα-activated activity (set at 100 %). n = 3-9.

Under identical experimental conditions 10 μM SDPA significantly potentiated mAC7 (2.4-fold), mAC9 (2.1-fold) and mAC6 activities (1.5-fold). Concentration-response curves were carried for mACs 1, 2, 6, 7 and 9 (Appendix Figure 3). The EC_50_ concentrations of SDPA for mAC6, 7 and 9 were 0.7 μM, i.e. not significantly different from suggesting equal binding affinities. The other Gsα-activated mAC isoforms were not significantly affected (Fig. 5 and Appendix Figure 3). In summary, the data demonstrated that the mAC isoform specificity of SDPA was not absolutely stringent. The data then pose the question whether other GPLs may exert similar effects on mAC activities or display a different panel of isoform specificity. This was investigated using four more stearoyl-2-docosahexaenoyl-GPLs (Fig. 5). 10 μM SDPS potentiated mAC3 and mAC9.

Smaller, yet significant effects were measured with mACs 7, 8, 5, and 6 (Fig. 5). 10 μM SDPE significantly potentiated Gsα-activated mAC isoforms 9, 3, 7, 5, and 4 (in this order). 10 μM SDPG significantly enhanced only mAC3 activity (Fig. 5). Compared to the seven-fold effect of SDPA on mAC3 these effects were small, yet, in mammalian biology such enhancements in mAC activity may well have profound physiological consequences. Up to 20 μM SDPC which is a major constituent of the outer leaflet of membranes had no effect on any mAC isoform (not shown). Taken together, the data then demonstrate the capacity of chemically defined GPLs to enhance or potentiate the activation of Gsα-activated mACs. We can virtually exclude coincidental and unspecific effects of the amphiphilic phospholipids because mAC isoforms were affected differentially. The results strongly suggest that a defined conformational space must exist at mACs which allows specific interactions with GPLs. Presently, the molecular details of the binding mode remain unknown.

### Relationship between SDPA and forskolin

SDPA failed to activate basal mAC3 activity and only potentiates Gsα-activated mAC3 activity (Fig. 1). The plant diterpene forskolin stimulates basal as well as Gsα-activated mAC activities (Dessauer et al., 2017, Tesmer, Sunahara et al., 1997, Zhang, Liu et al., 1997), i.e. the effects of SDPA and forskolin are only partly similar. Forskolin stimulates mACs expressed in Sf9 cells to rather different extents and with discrepant potencies, e.g. the EC_50_ concentrations for AC1 (0.7 μM) and AC2 (8.7 μM) differ more than 12-fold (Pinto, Papa et al., 2008). We established forskolin concentration-response curves for all mAC isoforms expressed in HEK293 cells under identical experimental conditions using Mg^2+^ as divalent cation, a comprehensive study which is lacking so far (Appendix Figure 4).

Stimulations at 1 mM forskolin were between 3-fold for mAC1 and remarkable 42-fold for mAC3. The EC_50_ concentrations ranged from 2 μM (mAC1) to 512 μM forskolin (mAC7; Appendix Figure 4). We also observed forskolin activation of mAC9 although the current consensus regarding this isoform is that it is forskolin insensitive. The latter conclusion is based on experiments with mAC9 expressed in insect Sf9 cells using Mn^2+^ as a cation (Dessauer et al., 2017). Another report described forskolin activation of mAC9 when expressed in HEK293 (Premont, Matsuoka et al., 1996), in line with our data (Appendix Figure 4). We examined potential interactions between forskolin and SDPA using mAC3 activated by 600 nM Gsα. Up to 10 μM, SDPA did not significantly affect forskolin stimulation. We reason that the absence of interactions or cooperativity between forskolin and SDPA suggests that both agents affect mAC regions which exclude mutual cooperative interactions. Nevertheless, considering the structural dissimilarity of forskolin and SDPA and the obvious lack of a molecular fit an identical binding site for both lipophilic agents is rather unlikely. On the other hand, both agents do interact with distantly-binding Gsα in a cooperative manner.

### SDPA enhances Gsα-stimulated cAMP formation in mouse brain cortical membranes

Above we tested GPLs with individual mAC isoforms. At this point the question is whether SDPA would potentiate mAC activity in membranes isolated from mammalian organs. Several mAC isoforms are expressed in any given tissue and cell, yet the ratios of isoform expression differ. In mouse brain cortex all mAC isoforms with the exception of mAC4 are expressed (Ludwig & Seuwen, 2002, Sanabra & Mengod, 2011). Depending on the expression ratios we might expect at least a moderate potentiation of the Gsα-activated AC activity by SDPA. In mouse cortical membranes the basal AC activity of 0.3 nmoles cAMP·mg^-1^·min^-1^ was unaffected by 10 μM SDPA (Fig. 6A). 600 nM Gsα stimulated AC activity 20-fold (7.9 nmoles cAMP·mg^-1^·min^-1^) and this was further enhanced 1.7-fold by 10 μM SDPA (13.4 nmoles cAMP·mg^-1^·min^-1^). An SDPA concentration-response curve yielded an EC_50_ of 1.2 μM, i.e. similar to those established in HEK293-expressed mAC isoforms (Fig. 6A; compare to Fig. 1 and Appendix Figure 3). The data demonstrated that the SDPA effect of GPLs on mAC activities was not due to peculiar membrane properties of the cultured HEK293 cells and supported the suggestion that the effects of GPLs are of more general physiological relevance.

**Figure 6.**
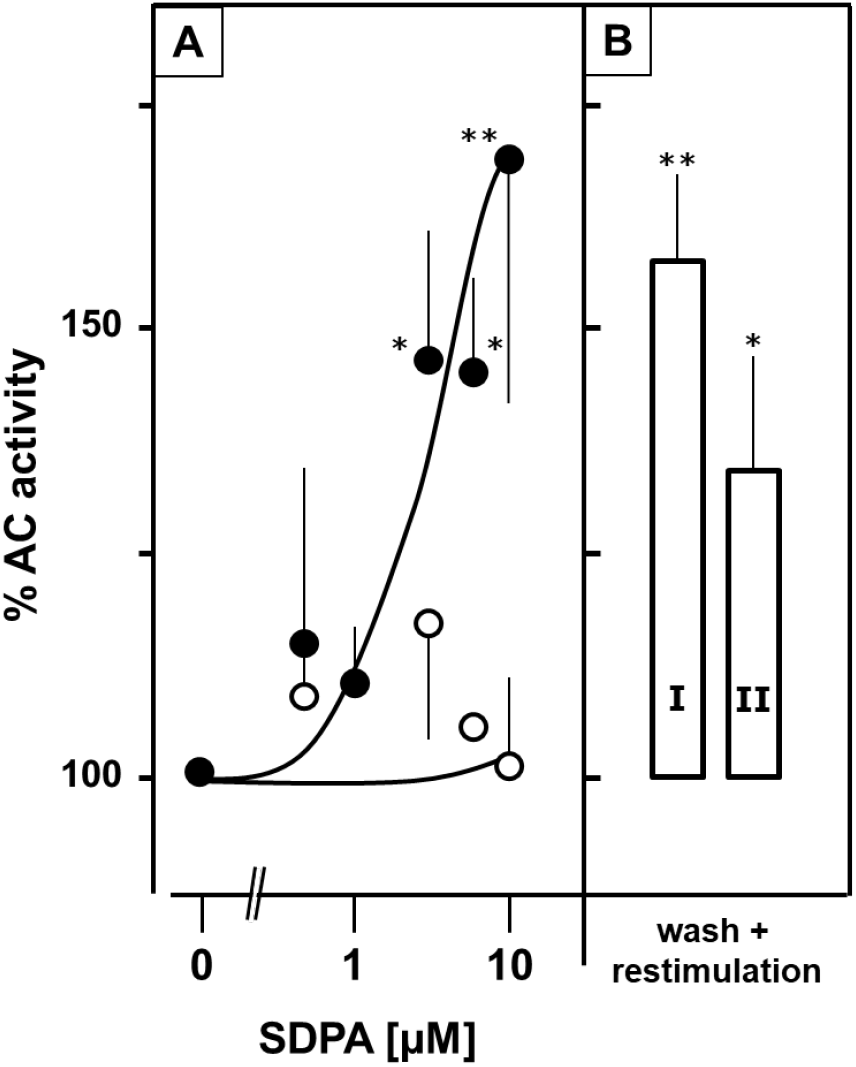
1-Stearoyl-2-docosahexaenoyl-phosphatidic acid (SDPA) concentration-dependently potentiates Gsα activated adenylyl cyclase activity in brain cortical membranes from mouse. (A) 600 nM Gsα was used to activate mACs in cortical membranes (solid circles: 100 % Gsα-activated activity is 7.9 ± 1.9 nmol cAMP•mg^-1^•min^-1^); open circles: basal activity (in absence of Gsα) is 0.3 ± 0.2 nmol cAMP•mg^-1^•min^-1^. n = 6. (B) Reversibility of SDPA action. Cortical brain membranes were incubated for 15 min without (I) and with (II) 10 μM SDPA. Membranes were then collected by centrifugation, and re-assayed + 600 nM Gsα and 10 μM SDPA. Error bars denote S.E.M., *: p< 0.05; **: p< 0.01 (n=6).

The next question was whether SDPA acts directly from the extracellular site via the membrane-receptor domain of hAC3 or is a cytosolic effector. We used HEK293 cells transfected with hAC3. 2.5 μM isoproterenol increased cAMP levels from 0.06 to 0.24 pmol/10^4^ cells within 45 min (6-fold stimulation). Including 10 μM SDPA in the medium did not affect isoproterenol stimulation. Similarly, we used 10 μM adenosine and 1 μM prostaglandin E_2_. Addition of SDPA did not enhance intracellular cAMP generation (see Appendix Table 3). These unequivocal data virtually excluded that SDPA acted via extracellular binding sites (receptors) or via an efficient and rapid uptake system. In fact, although SDPA is a GPL, it is unlikely that it can pass into intact cells. First, SDPA has a negatively charged headgroup which at the physiological pH of incubations is dissociated. Second, SDPA might slide into the outer leaflet of the membrane, but it would require a flippase for incorporation into the inner leaflet and potential release into the cytosol. Considering that the inner leaflet of the plasma membrane usually has a negative surface charge due to the predominance of phosphatidylserine we consider this as unlikely. Third, we did not observe significant incorporation of SDPA into brain cortical membranes (see below). Thus, the data tentatively suggest a cytosolic site for the action of GPLs.

### Is SDPA a ligand?

GPLs are common building blocks of cell membranes. Major constituents of the inner leaflet are phosphatidylserines and phosphatidylethanolamines, whereas the predominant lipids of the outer leaflet are phosphatidylcholine and sphingomyelin. In many tissues docosahexaenoic acid is a major acyl substituent in membrane GPLs (Hishikawa, Valentine et al., 2017). Phosphatidic acids are indispensable, yet minor membrane components (Kooijman & Burger, 2009). The potentiation by SDPA of Gsα-activated mAC3 could be due to a lack of SDPA in the vicinity of the membrane-imbedded mACs. Added SDPA might be incorporated into the membrane or inserted into hydrophobic pockets close to mACs resulting in an irreversible reordering of mAC domains. Alternatively, SDPA may bind reversibly to the cyclase in a transient manner. Under these latter circumstances the biochemical effect should be reversible. Using mouse cortical brain membranes, we attempted to dissect these possibilities. We incubated membranes for 15 min at 37°C with 10 μM SDPA. The membranes were then collected at 100,000 g and washed once. The pre-treated membranes were susceptible to Gsα stimulation and concomitant potentiation by SDPA like naïve cortical membranes (Fig. 6B). Furthermore, the supernatant of a 50 μM SDPA preincubation was used to potentiate the Gsα activation in naïve membranes, i.e. SDPA was not significantly incorporated into the membrane preparation. The data support the notion that SDPA, and most likely other GPLs, serve as intracellular effectors for mACs.

## Discussion

Our results were contrary to the hypothesis at the outset because we expected to find an mAC inhibitory input. Most surprisingly we identified SDPA and other GPLs as positive effectors of mAC activities. Obviously, we have discovered a new system of intracellular mAC regulation. At this state our findings open more questions than can be answered with this initial report.

We used HEK293 cells permanently transfected with mACs. HEK293 cells express considerable endogenous AC3 and 6 activities (Soto-Velasquez, Hayes et al., 2018). These endogenous mAC activities appear to be negligible in this context. First, upon transfection of mAC isoforms we observed very different basal AC activities virtually excluding that ‘contaminating’ endogenous AC activities affected our results (see Appendix Table 2 for a list of basal activities in transfected HEK293 cells). Second, we tested HEK293 cells in which mACs 3 and 6 were knocked out (Soto-Velasquez et al., 2018). Upon mAC3 transfection SDPA similarly potentiated Gsα-activated mAC3 activity. Because these engineered cells proliferated rather slowly they were not used routinely.

Diacylglycerols’ and PA are lipid second messengers that regulate physiological and pathological processes, e.g. phosphatidic acids were reported to effect ion channel regulation and SDPA to act on the serotonin transporter in the brain (Lu, Murakami et al., 2020, Robinson, Rohacs et al., 2019, Shin & Loewen, 2011). So far, the specificity of fatty acyl residues and head groups in these lipids was not explored. Here, we observed a striking exclusivity of fatty-acyl esters at positions 1- and 2-of the glycerol backbone supporting a specific effector-mAC interaction. Usually fatty acyl substitutions are regulated because they impart specific biophysical and biochemical properties (Hishikawa et al., 2017). We demonstrated that the combined fatty acyl ligands 1-stearoyl-2-docosahexaenoyl are more or less exclusive for the actions of SDPA. Even seemingly minor changes caused substantial changes in activity and efficacy, e.g. a change from stearoyl to palmitoyl at glycerol position 1 (Fig. 3). This argues for a specific steric interaction between the flexible stearoyl- and docosahexaenoyl carbon-chains. Acyl chain substitutions might then impair specific protein-ligand interactions, e.g. by a shrinkage of the binding surface. Apparently, such interactions are substantially diminished when only one of the two acyl residues is altered. Notably, di-docosahexaenoyl- and di-arachidonoyl-phosphatidic acids (DDPA and DAPA) had mostly lost the capability to promote AC3 activity (Fig. 3). A particularly interesting point is the preference for 2-docosahexaenoyl acylation in the GPLs. Docosahexaenoic acid is an essential omega-3 fatty acid that cannot be synthesized at adequate quantities in infants or seniors (Qiu, 2003). Therefore it is widely sold as a nutraceutical and should be included into a balanced diet. Docosahexaenoic acid is particularly abundant in membrane lipids in the retina (about 60 % of all lipids contain docosahexaenoic acid), testes, brain, heart and skeletal muscle (Hishikawa et al., 2017) and a sodium-dependent symporter for uptake of this fatty acid, packaged as a lysophosphatidic acid, has been characterized and its structure was elucidated by cryo-EM (Cater, Chua et al., 2021, Nguyen, Ma et al., 2014, Wood, Zhang et al., 2021). Docosahexaenoic acid is needed for normal brain development and cognitive functions, a role in depression, aging and Alzheimer’s disease is discussed (Duan, Song et al., 2021, Heras-Sandoval, Pedraza-Chaverri et al., 2016, Hishikawa et al., 2017, Nguyen et al., 2014, Zhu, Tan et al., 2015). So far, mACs have not yet been noticed in metabolic disturbances caused by a lack of docosahexaenoic acid. The data presented here provides evidence that docosahexaenoic acid is involved in stimulating the cAMP generating system.

Examination of head group specificity displayed different patterns of mAC susceptibility and activity (Fig. 5). Notably, mAC isoforms 1 and 2 were not significantly affected by any of the GPLs assayed. This may be due to a general insensitivity for GPLs or that we did not identify the suitable bioactive GPLs. We did not examine the specificity of acyl substitution at the glycerol backbone in SDPS, SDPE, SDPG and SDPC because of the specificity of the stearic/docosahexaenoic acid couple in SDPA. We tested 1-stearoyl-2-arachidonoyl-phosphatidyl choline and the corresponding phosphatidyl-ethanolamine. Biological activity was absent with mACs 3, 5, 7, and 9, bolstering the assertion that fatty acyl specificity is stringent in these GPLs as well. Presently we cannot completely exclude that GPLs acylated by different couples of acid substituents at the 1- and 2-positions might possess equal or better effector properties. In view of the large variety of GPLs this cannot be tested with a reasonable effort. Currently, we consider such a possibility as remote. We do not know how GPL’s mechanistically potentiate AC activity in a synergistic interaction together with Gsα. The tentative scheme in Fig. 7 is intended to illustrate an approximation of potential interaction sites in relation to Gsα and forskolin. The precise nature of such interactions requires structural details (in progress).

**Figure 7.**
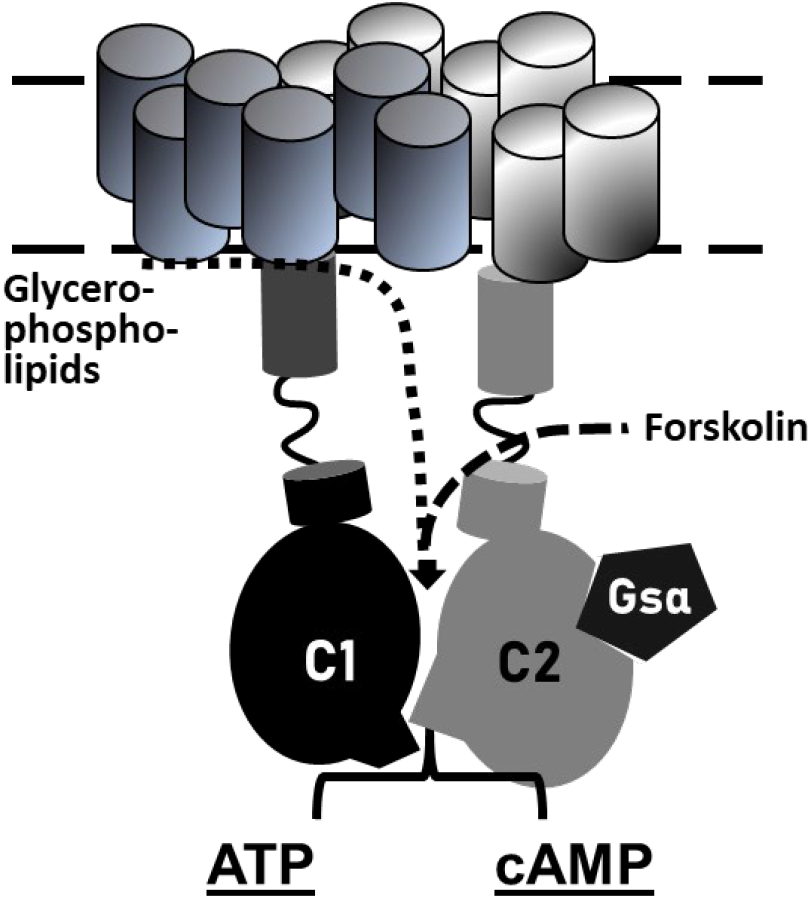
Tentative scheme of a 2X6TM-adenylyl cyclase with regulatory input from Gsα, binding to the C2 catalytic domain, forskolin, binding to a degenerated second substrate-binding site (Guo et al., 2001, Tesmer & Sprang, 1998), and glycerophospholipids, here proposed to enter and bind at the membrane anchor-receptor and extending towards the catalytic dimer.

Another question which is not answered in this study concerns the intracellular origin of GPLs, how their biosynthesis and release is regulated and tied into the cAMP regulatory system. Despite being water insoluble, an efficient traffic of phospholipids in cells exists, e.g. between locations of uptake and biosynthesis, to and from low-density-, high-density- and very low density lipoproteins, and the diversity of membrane-enclosed organelles such as mitochondria, nucleus, endoplasmic reticulum, endosomes, lysosomes, and the plasma membrane itself. Thus, lipid trafficking is a continuous cellular process connected to diverse signaling systems (Hishikawa et al., 2017, Shin & Loewen, 2011). Part of the biosynthetic pathways for phosphatidic acid is the hydrolysis of GPLs with choline, ethanolamine or serine as headgroups by phospholipase D which generates phosphatidic acids (Jang, Lee et al., 2012). Chemically, GPLs are excellently suited to serve as mAC effectors because termination of SDPA signaling is easily accomplished by phospholipase C. The relationship between SDPA and forskolin is debatable. The agents do not cooperatively interact at mAC proteins. Certainly, the structural changes caused by either agent promote the interactions between AC and Gsα. Yet this is no proof that such changes are identical or even similar.

A critical observation was the potentiation by SDPA of Gsα-activated mAC activity in mouse brain cortical membranes. mAC3 has been reported to be abundantly expressed in brain (Ludwig & Seuwen, 2002, Sanabra & Mengod, 2011). The efficacy of SDPA was comparable to that determined in mAC3-HEK293 membranes. This demonstrated that the effect of GPLs observed in HEK293 expressed AC isoforms is of physiological significance. Our approach has then discovered intracellular processes which in conjunction with the established canonical GPCR/Gsα-regulation of mACs add a new dimension of mAC regulation. Currently, we cannot exclude the possibility that other GPLs exist which have an inhibitory input. Actually, thermodynamic considerations would argue in favor of such a possibility. Whether this is realized as a biological mechanism remains an open possibility. Presently, many important questions remain unanswered, such as how are intracellular GPL levels regulated, which of the intracellular GPLs have access to the membrane-delimited ACs, are GPL concentrations persistently or acutely adjusted in a cell, e.g. by stress, diet, diurnal or seasonal effects or by peculiar disease states? In other word, are we dealing with a long-term regulation of the Gsα-sensitivity of the cAMP generating system or with coordinated short term signaling events? Answering these medically relevant questions remains a formidable challenge in the future.

## Materials and methods

The genes of the human AC isoforms 1 – 9 cloned into the expression plasmid pcDNA3.1+/C-(K)-DYK were purchased from GenScript and contained a C-terminal flag-tag. Creatine kinase was purchased from Sigma, restriction enzymes from New England Biolabs or Roche Molecular. All chemicals were from Avanti Lipids and Sigma-Merck. The constitutively active GsαQ227L point mutant was expressed and purified as described earlier (Diel, Klass et al., 2006, Graziano, Freissmuth et al., 1989, Graziano et al., 1991). Forskolin was a gift from Hoechst, Frankfurt, Germany. Human serum (catalog # 4522 from human male AB plasma) and fetal bovine serum were from Gibco, Life Technologies, Darmstadt, Germany (catalog #: 10270; lot number: 42Q8269K).

### Plasmid construction and Protein Expression

ACIC1_ACIIC2 was generated in pQE60 with NcoI/HindIII restrictions sites according to Tang et al. (Tang & Gilman, 1995). The construct boundaries were: MRGSH_6_-HA-hAC1-C1_M268–R482_-AAAGGMPPAAAGGM -hAC2-C2_R822-S1091_. HEK293 cells were maintained in Dulbecco’s modified Eagle’s medium (DMEM) containing 10% fetal bovine serum at 37°C with 5% CO2. Transfection of HEK293 cells with single mAC plasmids was with PolyJet (SignaGen, Frederick, MD, USA). Permanent cell lines were generated by selection for 7 days with G418 (600 μg/mL) and maintained with 300 μg/mL G418 (Baldwin, Li et al., 2019, Cumbay & Watts, 2004, Soto-Velasquez et al., 2018). For membrane preparation cells were tyrpsinized and collected by centrifugation (3,000×g, 5 min). Cells were lysed and homogenized in 20 mM HEPES, pH 7.5, 1 mM EDTA, 2 mM MgCl_2_, 1 mM DTT, and one tablet of cOmplete, EDTA-free (for 50 mL), 250 mM sucrose by 20 strokes in a potter homogenizer. Debris was removed by centrifugation for 5 min at 1,000 × g, membranes were then collected by centrifugation at 100,000 × g for 60 min at 0°C, resuspended and stored at -80°C in 20 mM MOPS, pH 7.5, 0.5 mM EDTA, 2 mM MgCl_2_. Expression was checked by Western blotting.

Membrane preparation from mouse brain cortex was according to (Schultz & Schmidt, 1987, Seth et al., 2020). For each preparation three cerebral cortices were dissected and homogenized in 4.5 ml cold 48 mM Tris-HCl, pH 7.4, 12mM MgC1_2_, and 0.1 mM EGTA with a Polytron hand disperser (Kinematica AG, Switzerland). The homogenate was centrifuged for 15min at 12,000g at 4°C and the pellet was washed once with 5 mL 1 mM potassium bicarbonate. The final suspension in 2 mL 1 mM KHCO_3_ was stored in aliquots at −80°C.

### Adenylyl cyclase assay

AC activities were determined in a volume of 10 μl using 1 mM ATP, 2 mM MgCl_2_, 3 mM creatine phosphate, 60 μg/ml creatine kinase, 50 mM MOPS, pH 7.5 using the cAMP assay kit from Cisbio (Codolet, France) according to the supplier’s instructions. For each assay a cAMP standard curve was established (Seth et al., 2020). Lipids were dissolved in 100% ethanol or DMSO at high concentrations and acutely diluted in 20 mM MOPS pH 7.5 at concentrations which limited organic solvent in the assay at maximally 1%. Up to 2 % neither ethanol nor DMSO had any effect on AC activities.

### Lipidomic analysis

Lipids were extracted from MonoQ purified aqueous fractions by methyl-*tert*-butyl ether / methanol as described (Matyash et al., 2008) after adjusting their pH to 1.0 and 6.0, respectively. The collected extracts were dried under vacuum, and re-dissolved in 500μl of water / acetonitrile 1:1 (v/v). Lipids were analyzed by LC-MS/MS on a Xevo G2-S QTof (Waters) mass spectrometer interfaced to Agilent 1200 liquid chromatograph. Lipids were separated on a Cortecs C18 2.7 μm beads; 2.1 mm ID x 100 mm (Waters) using a mobile phase gradient: solvent A: 50% aqueous acetonitrile; solvent B: 25% of acetonitrile in isopropanol; both A and B contained 0.1% formic acid (v/v) and 10 mM ammonium formate. The linear gradient was delivered with flow rate of 300 μl /min in 0 min to 12 min from 20% to 100 % B; from 12 min to 17 min maintained at 100% B, and from 17 min to 25min at 20% B. Mass spectra were acquired within the range of *m/z* 50 to m/z 1200 at the mass resolution of 20 000 (FWHM). The chromatogram was searched against web-accessible XCMS compound database at https://xcmsonline.scripps.edu/landing_page.php?pgcontent=mainPage. Lipids were quantified using Skyline 21.1.0.278 software using synthetic lipid standards (Vvedenskaya et al., 2021) spiked into the analyzed fractions prior lipid extraction.

### Data analysis and statistical analysis

All incubations were in duplicates or triplicates. For easier presentations data were normalized to respective controls and n and S.E.M values are indicated in all figures. Data analysis was with GraphPad prism 8.1.2 using a two-tailed t-test.

## Abbreviations used

mAC: membrane-delimited adenylyl cyclase
GPL: glycerophospholipid
SDPA: 1-stearoyl-2-docosahexaenoyl-phosphatidic acid

## Supplemental Material

**Appendix Figure 1.**
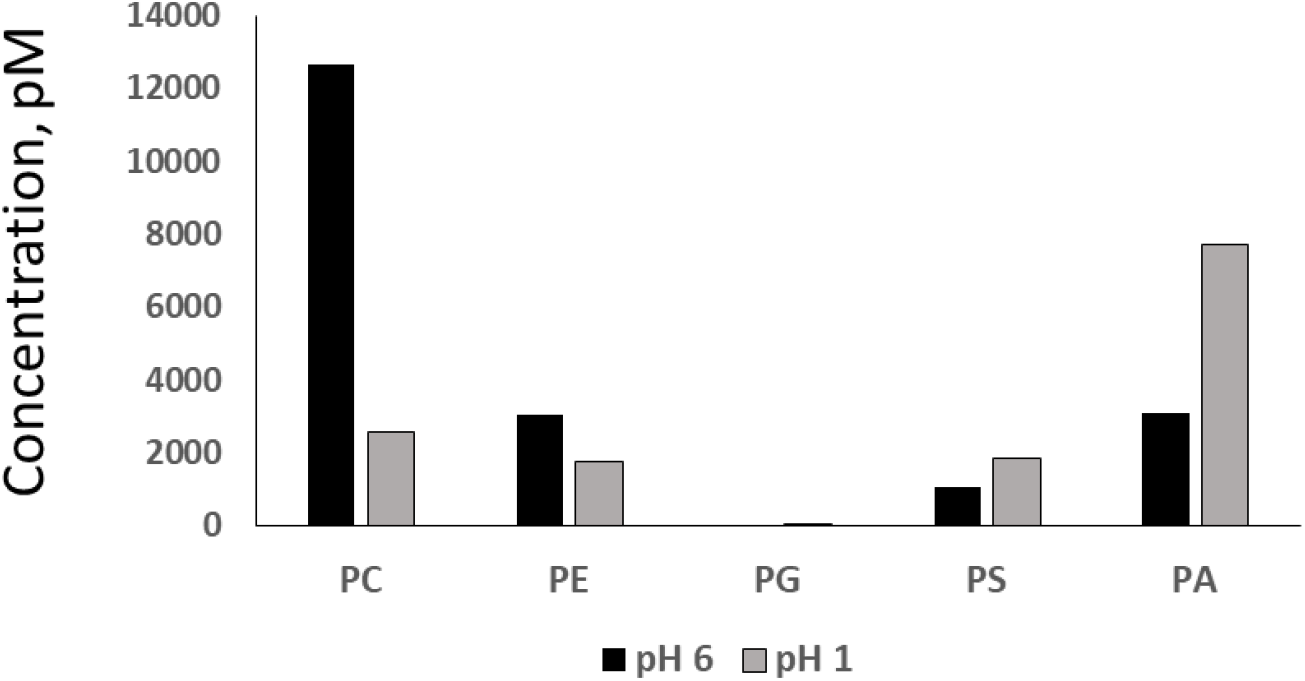
Lipid class composition of MTBE / methanol extracts. MonoQ-purified fractions were extracted at pH 1.0 and pH 6.0. Expectantly, the extract recovered under acidic conditions was enriched with PA. Y-axis: total abundance of lipid classes, pmol/L (n=2).

**Appendix Figure 2.**
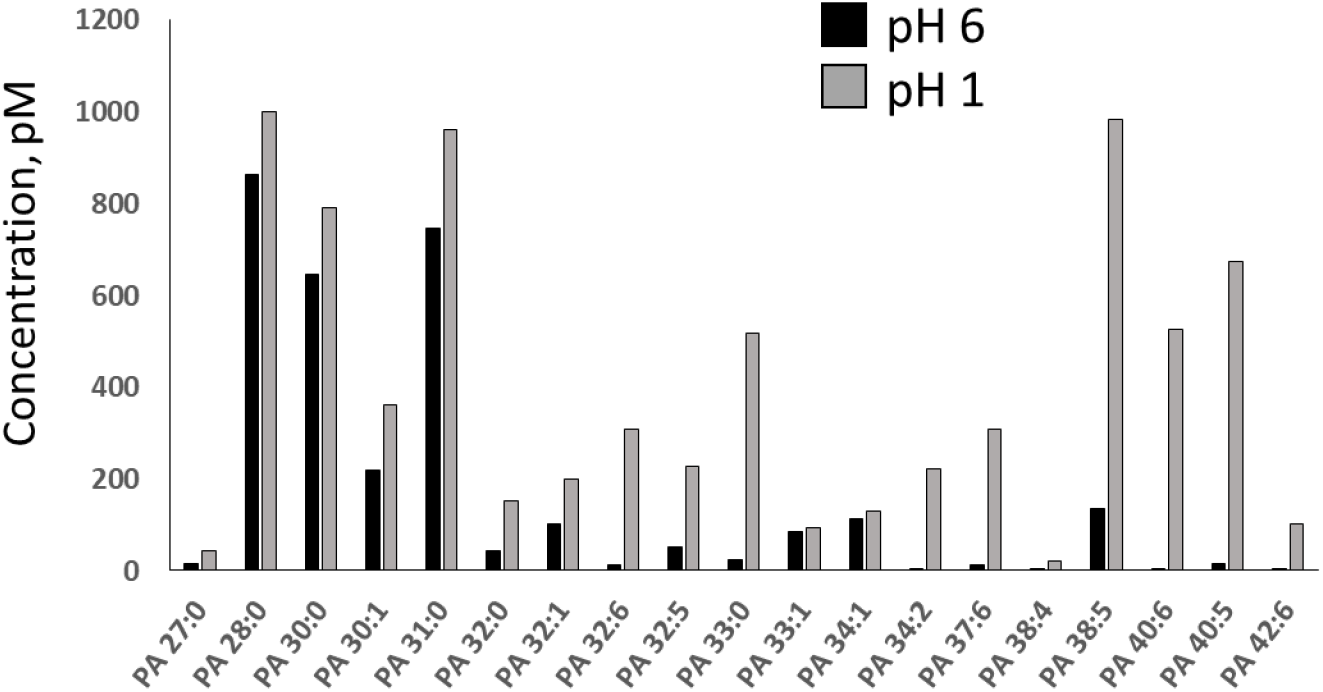
Molecular composition of PA species extracted by MTBE / methanol from the fractions with pH 6.0 and pH 1.0. Acidic extraction increased the recovery of PA by more than 2-fold and also enriched the extract with the molecular species comprising long polyunsaturated fatty acid moieties. Y-axes: molar abundance of lipid species, in pmol/L (n=2).

**Appendix Table 1:** List of lipids tested: from Avanti lipids:

- 131303P Cerebrosides
- 131305P Sulfatides
- 800818C-1-stearoyl-2-arachidonoyl-sn-glycerol
- 800819 --stearoyl-2-docosahexaenoyl-sn-glycerol
- 830855C 1,2-dipalmitoyl-sn-glycero-3-phosphate
- 840051P L-α-phosphatidylcholine (Egg, Chicken)
- 840055C L-α-phosphatidylcholine (Liver, Bovine)
- 840065C 1-stearoyl-2-docosahexaenoyl-sn-glycero-3-phospho-L-serine
- 840101C L-α-phosphatidic acid (Egg, Chicken) (sodium salt)
- 840859C 1-palmitoyl-2-arachidonoyl-sn-glycero-3-phosphate (sodium salt)
- 840860C 1-palmitoyl-2-docosahexaenoyl-sn-glycero-3-phosphate (sodium salt)
- 840862C 1-stearoyl-2-linoleoyl-sn-glycero-3-phosphate (sodium salt)
- 840863C 1-stearoyl-2-arachidonoyl-sn-glycero-3-phosphate (sodium salt)
- 840864C 1-stearoyl-2-docosahexaenoyl-sn-glycero-3-phosphate (sodium salt)
- 840875C 1,2-dioleoyl-sn-glycero-3-phosphate (sodium salt)
- 840885C 1,2-dilinoleoyl-sn-glycero-3-phosphate (sodium salt)
- 840886C 1,2-diarachidonoyl-sn-glycero-3-phosphate (sodium salt)
- 840887C 1,2-didocosahexaenoyl-sn-glycero-3-phosphate (sodium salt)
- 850469C 1-stearoyl-2-arachidonoyl-sn-glycero-3-phosphocholine
- 850472C 1-stearoyl-2-docosahexaenoyl-sn-glycero-3-phosphocholine
- 850804C 1-stearoyl-2-arachidonoyl-sn-glycero-3-phosphoethanolamine
- 850806C 1-stearoyl-2-docosahexaenoyl-sn-glycero-3-phosphoethanolamine
- 850852C 1,2-dioleoyl-sn-glycero-3-phosphoethanolamine-N,N-dimethyl
- 857130P 1-oleoyl-2-hydroxy-sn-glycero-3-phosphate (sodium salt)
- 857328P 1-oleoyl-sn-glycero-2,3-cyclic-phosphate (ammonium salt)
- 860053P total ganglioside extract (Brain, Porcine-Ammonium Salt)
- 860492 Sphingosine-1-phosphate; D-erythro-sphingosine-1-phosphate LIPOID (Heidelberg) donated the following lipids:
- 30. 556200 Lipoid PC 14:0/14:0; 1,2-Dimyristoyl-sn-glycero-3-phosphatidylcholine (DMPC)
- 31. 556300 Lipoid PC 16:0/16:0;1,2-Dipalmitoyl-sn-glycero-3-phosphatidylcholine (DPPC)
- 32. 556500 Lipoid PC 18:0/18:0; 1,2-Distearoyl-sn-glycero-3-phosphocholine (DSPC)
- 33. 556600 Lipoid PC 18:1/18:1; 1,2-Dioleoyl-sn-glycero-3-phosphocholine (DOPC)
- 34. 556400 Lipoid PC 16:0/18:1; 1-Palmitoyl-2-oleoyl-sn-glycero-3-phosphocholine (POPC)
- 35. 557100 Lipoid PC 22:1/22:1; 1,2-Dierucoyl-sn-glycero-3-phosphocholine (DEPC)
- 36. 566300 Lipoid PA 16:0/16:0; 1,2-Dipalmitoyl-sn-glycero-3-phosphate, mono-sodium salt (DPPA-Na)
- 37. 567600 Lipoid PS 18:1/18:1; 1,2-Dioleoyl-sn-glycero-3-phosphoserine, sodium salt (DOPS-Na)
- 38. 560200 Lipoid PG 14:0/14:0; 1,2-Dimyristoyl-sn-glycero-3-phospho-rac-glycerol-Na (DMPG)
- 39. 560300 Lipoid PG 16:0/16:0; 1,2-Dipalmitoyl-sn-glycero-3-phospho-rac-glycerol-Na (DPPG)
- 40. 560400 Lipoid PG 18:0/18:0; 1,2-Distearoyl-sn-glycero-3-phospho-rac-glycerol-Na (DSPG)
- 41. 565600 Lipoid PE 14:0/14:0; 1,2-Dimyristoyl-sn-glycero-3-phosphoethanolamine (DMPE)
- 42. 565300 Lipoid PE 16:0/16:0; 1,2-Dipalmitoyl-sn-glycero-3-phosphoethanolamine (DPPE)
- 43. 565400 Lipoid PE 18:0/18:0; 1,2-Distearoyl-sn-glycero-3-phosphoethanolamine (DSPE)
- 44. 565600 Lipoid PE 18:1/18:1; 1,2-Dioleoyl-sn-glycero-3-phosphoethanolamine (DOPE)

**Appendix Table 2.**
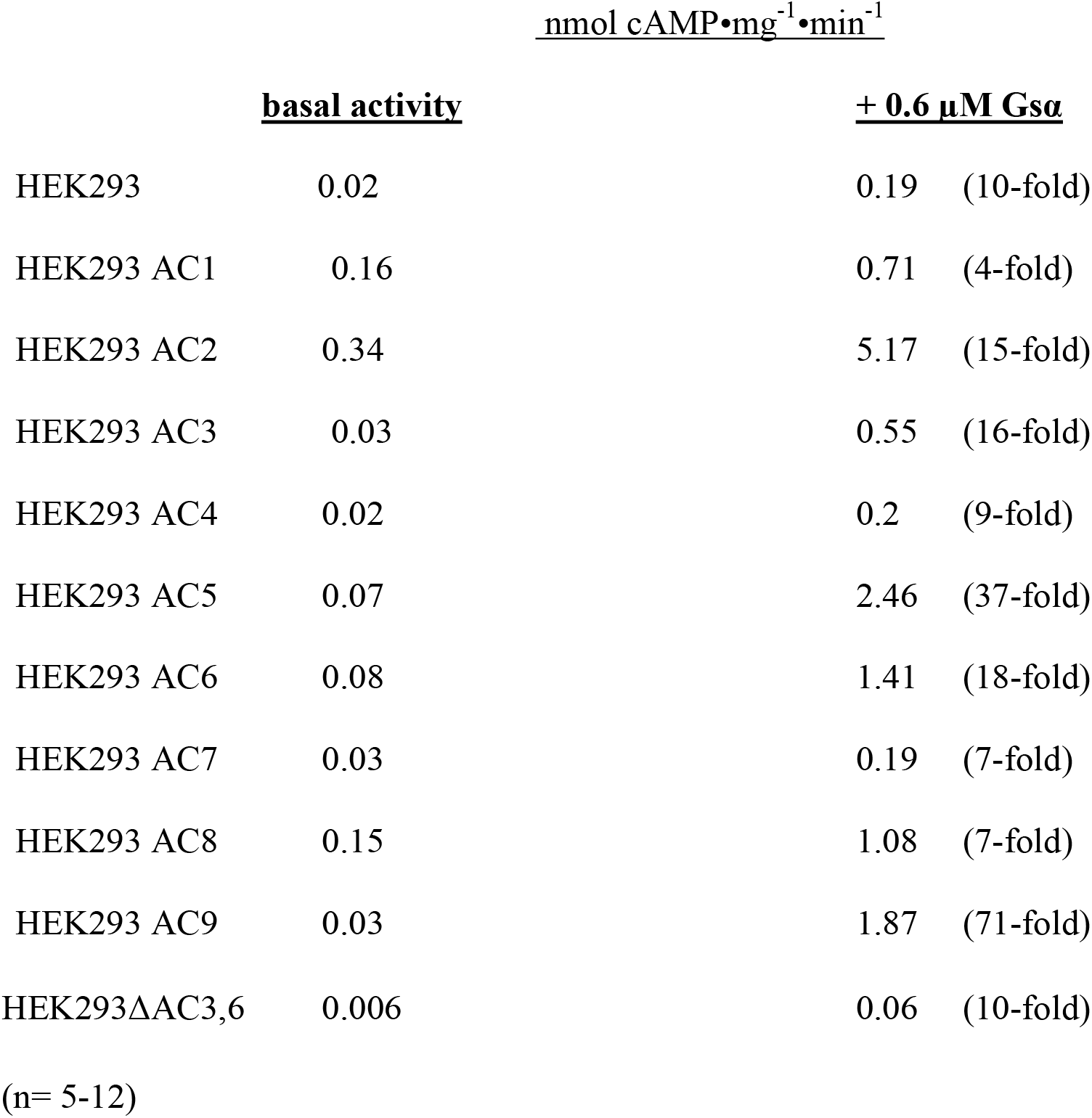
mAC activities in HEK293 cell membranes transfected with human mAC isoforms

**Appendix Figure 3.**
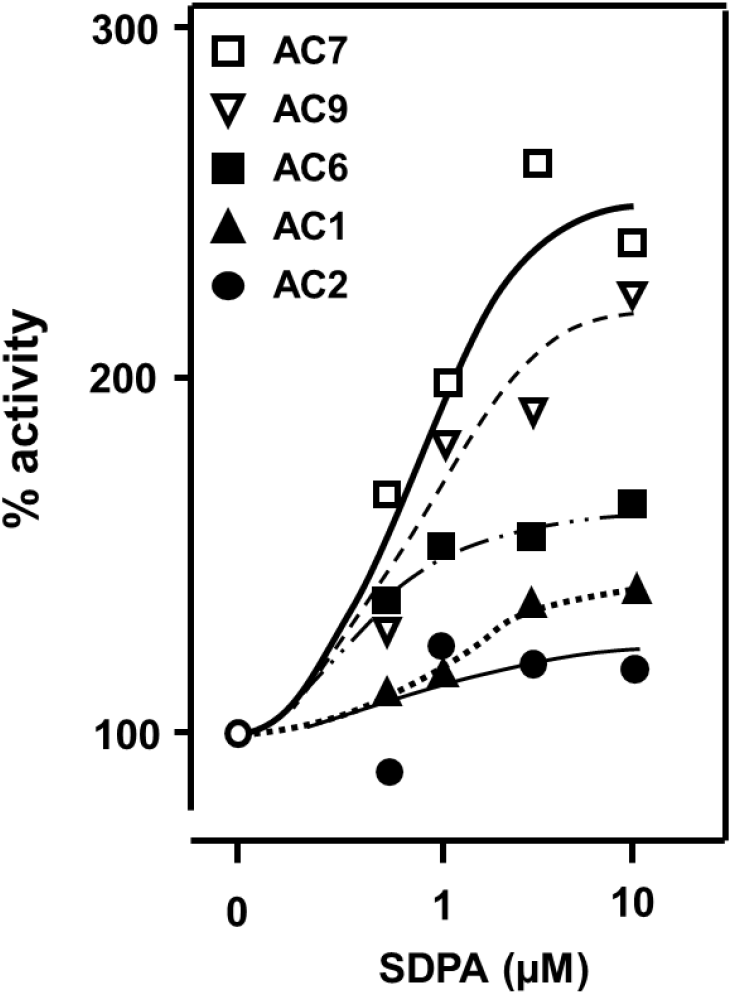
Concentration-response curves for SDPA potentiation of mAC isoforms 7, 9, 6, 1, and 2. Basal and Gsα-activated activities are listed in Appendix table 2. n=2-5.

**Appendix Figure 4:**
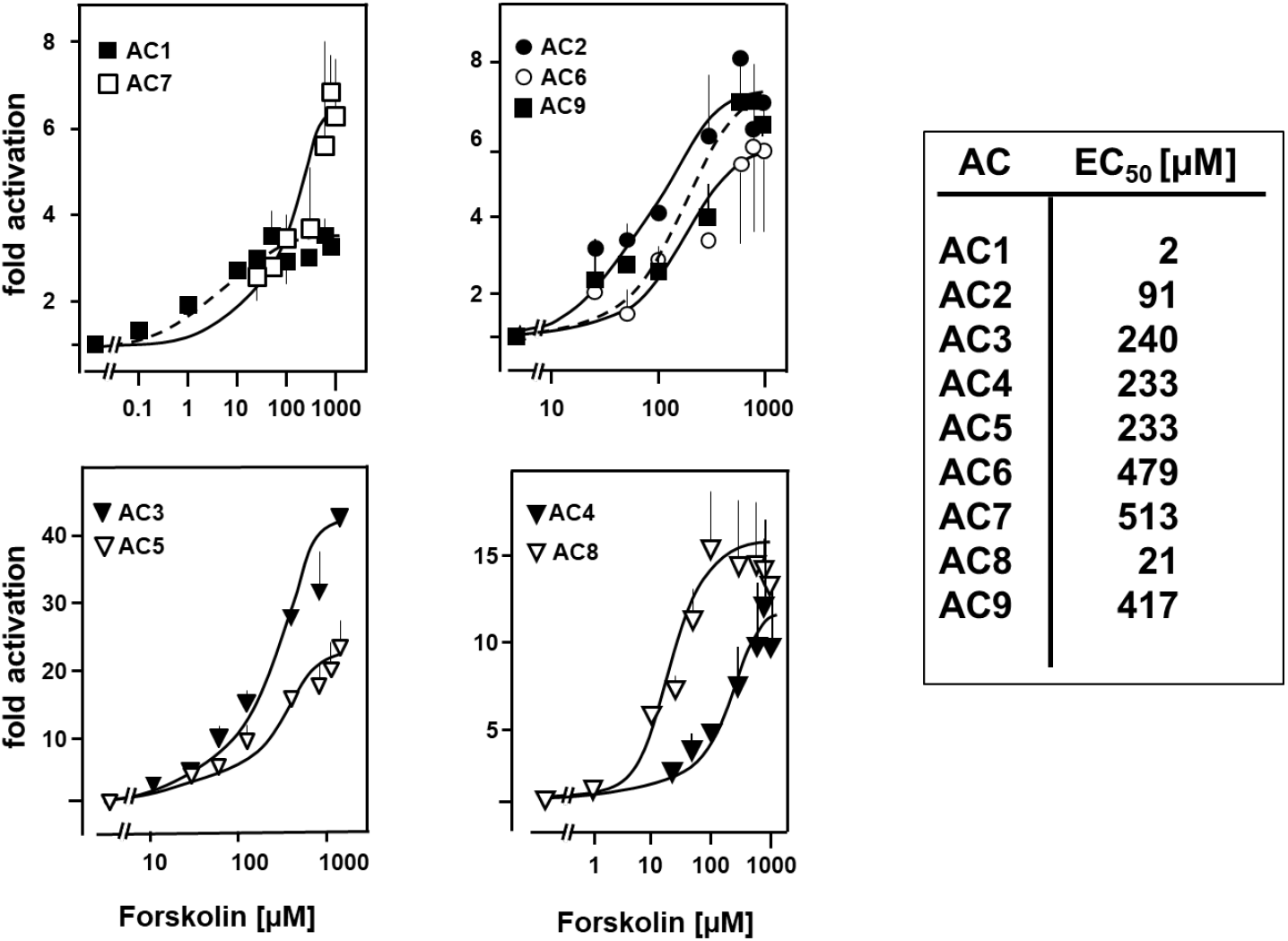
Forskolin concentration-response curves for the nine human mAC isoforms expressed in HEK293 cells. Error bars denote S.E.M. The calculated EC_50_ concentrations are listed at right. n = 2-4.

**Appendix Table 3.** With hAC3 transfected HEK293 cells in a 396 well plate were incubated and stimulated at 37°C for 45 min by adenosine, isoproterenol and prostaglandin E_2_ ± 10 μM SDPA. n = 3 to 4, mean ± S.E.M. Incubations were stopped by addition of detection and lysis buffer of the cAMP assay kit (10 μl/well; Cisbio).

|  | pMoles cAMP/<br>10<sup>4</sup> cells |
| --- | --- |
| <u>basal</u> | 0.06 ± 0.02 |
| <u>SDPA, 10 µM</u> | 0.02 ± 0.01 |
| <u>2.5 µM isoproterenol</u> | 0.24 ± 0.03 |
| <u>2.5 µM isoproterenol + 10 µM SDPA</u> | 0.25 ± 0.03 |
| <u>1 µM prostaglandin E<sub>2</sub></u> | 0.10 ± 0.01 |
| <u>1 µM prostaglandin E<sub>2</sub> + 10 µM SDPA</u> | 0.11 ± 0.02 |
| <u>10 µM adenosine</u> | 0.17 ± 0.07 |
| <u>10 µM adenosine + 10 µM SDPA</u> | 0.07 ± 0.01 |

## Acknowledgements

We thank U. Kurz for a continuous supply of Gsα (Q227L), Dr. V. Watts, for suppling us with HEK293ΔmAC3ΔmAC6 cells, and Prof. Dr. A. Lupas for continuous encouragement. We gratefully acknowledge constructive suggestions from Dr. J. Linder. Supported by the Deutsche Forschungsgemeinschaft and institutional funds from the Max-Planck-Society.

## Additional Information

### Funding

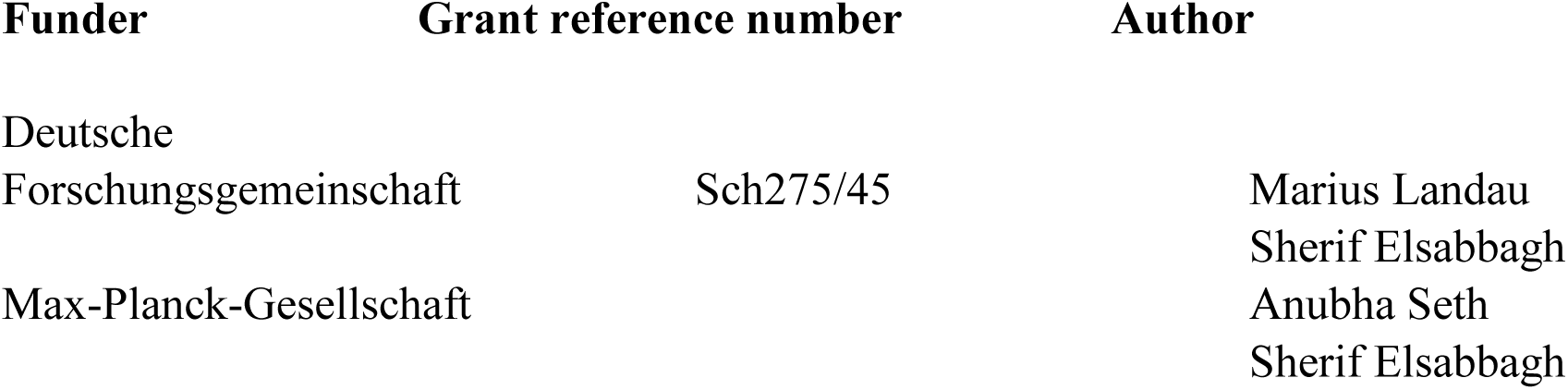

The funders had no role in study design, data collection and interpretation, or the decision to submit the work for publication.

### Author contributions

AS, ML, AS, acquisition of data, analysis and interpretation of data; SE, transfection and culture of permanent HEK293 cell lines with human adenylyl cyclases, AS, ST, lipidomic analysis and interpretation of data, JES, conception and design, analysis and interpretation of data, design of figures, writing manuscript.

## References

Baldwin TA, Li Y, Brand CS, Watts VJ, Dessauer CW (2019) Insights into the Regulatory Properties of Human Adenylyl Cyclase Type 9. Mol Pharmacol 95: 349–360

Bassler J, Schultz JE, Lupas AN (2018) Adenylate cyclases: Receivers, transducers, and generators of signals. Cell Signaling 46: 135–144

Beltz S, Bassler J, Schultz JE (2016) Regulation by the quorum sensor from Vibrio indicates a receptor function for the membrane anchors of adenylate cyclases. Elife 5

Bligh EG, Dyer WJ (1959) A rapid method of total lipid extraction and purification. Can J Biochem Physiol 37: 911–7

Cater RJ, Chua GL, Erramilli SK, Keener JE, Choy BC, Tokarz P, Chin CF, Quek DQY, Kloss B, Pepe JG, Parisi G, Wong BH, Clarke OB, Marty MT, Kossiakoff AA, Khelashvili G, Silver DL, Mancia F (2021) Structural basis of omega-3 fatty acid transport across the blood-brain barrier. Nature 595: 315–319

Cumbay MG, Watts VJ (2004) Novel regulatory properties of human type 9 adenylate cyclase. J Pharmacol Exp Ther 310: 108–15

Dessauer CW, Watts VJ, Ostrom RS, Conti M, Dove S, Seifert R (2017) International Union of Basic and Clinical Pharmacology. CI. Structures and Small Molecule Modulators of Mammalian Adenylyl Cyclases. Pharmacol Rev 69: 93–139

Diel S, Klass K, Wittig B, Kleuss C (2006) Gbetagamma activation site in adenylyl cyclase type II. Adenylyl cyclase type III is inhibited by Gbetagamma. J Biol Chem 281: 288–294

Duan J, Song Y, Zhang X, Wang C (2021) Effect of omega-3 Polyunsaturated Fatty Acids-Derived Bioactive Lipids on Metabolic Disorders. Front Physiol 12: 646491

Graziano MP, Freissmuth M, Gilman AG (1989) Expression of Gs alpha in Escherichia coli. Purification and properties of two forms of the protein. J Biol Chem 264: 409–18

Graziano MP, Freissmuth M, Gilman AG (1991) Purification of recombinant Gs alpha. Methods Enzymol 195: 192–202

Guo YL, Seebacher T, Kurz U, Linder JU, Schultz JE (2001) Adenylyl cyclase Rv1625c of Mycobacterium tuberculosis: a progenitor of mammalian adenylyl cyclases. EMBO J 20: 3667–3675

Heras-Sandoval D, Pedraza-Chaverri J, Perez-Rojas JM (2016) Role of docosahexaenoic acid in the modulation of glial cells in Alzheimer’s disease. J Neuroinflammation 13: 61

Hishikawa D, Valentine WJ, Iizuka-Hishikawa Y, Shindou H, Shimizu T (2017) Metabolism and functions of docosahexaenoic acid-containing membrane glycerophospholipids. FEBS Lett 591: 2730–2744

Jang JH, Lee CS, Hwang D, Ryu SH (2012) Understanding of the roles of phospholipase D and phosphatidic acid through their binding partners. Prog Lipid Res 51: 71–81

Kanacher T, Schultz A, Linder JU, Schultz JE (2002) A GAF-domain-regulated adenylyl cyclase from Anabaena is a self-activating cAMP switch. EMBO Journal 21: 3672–3680

Kooijman EE, Burger KN (2009) Biophysics and function of phosphatidic acid: a molecular perspective. Biochim Biophys Acta 1791: 881–8

Linder JU, Schultz JE (2003) The class III adenylyl cyclases: multi-purpose signalling modules. Cell Signal 15:1081–1089

Lu Q, Murakami C, Murakami Y, Hoshino F, Asami M, Usuki T, Sakai H, Sakane F (2020) 1-Stearoyl-2-docosahexaenoyl-phosphatidic acid interacts with and activates Praja-1, the E3 ubiquitin ligase acting on the serotonin transporter in the brain. FEBS Lett 594: 1787–1796

Ludwig MG, Seuwen K (2002) Characterization of the human adenylyl cyclase gene family: cDNA, gene structure, and tissue distribution of the nine isoforms. J Recept Signal Transduct Res 22: 79–110

Matyash V, Liebisch G, Kurzchalia TV, Shevchenko A, Schwudke D (2008) Lipid extraction by methyl-tert-butyl ether for high-throughput lipidomics. J Lipid Res 49: 1137–46

Ng WL, Wei Y, Perez LJ, Cong J, Long T, Koch M, Semmelhack MF, Wingreen NS, Bassler BL (2010) Probing bacterial transmembrane histidine kinase receptor-ligand interactions with natural and synthetic molecules. Proceedings of the National Academy of Sciences of the United States of America 107: 5575–80

Nguyen LN, Ma D, Shui G, Wong P, Cazenave-Gassiot A, Zhang X, Wenk MR, Goh EL, Silver DL (2014) Mfsd2a is a transporter for the essential omega-3 fatty acid docosahexaenoic acid. Nature 509: 503–6

Ostrom KF, LaVigne JE, Brust TF, Seifert R, Dessauer CW, Watts VJ, Ostrom RS (2022) Physiological roles of mammalian transmembrane adenylyl cyclase isoforms. Physiol Rev 102: 815–857

Pinto C, Papa D, Hubner M, Mou TC, Lushington GH, Seifert R (2008) Activation and inhibition of adenylyl cyclase isoforms by forskolin analogs. J Pharmacol Exp Ther 325: 27–36

Premont RT, Matsuoka I, Mattei MG, Pouille Y, Defer N, Hanoune J (1996) Identification and characterization of a widely expressed form of adenylyl cyclase. J Biol Chem 271: 13900–7

Qiu X (2003) Biosynthesis of docosahexaenoic acid (DHA, 22:6-4, 7,10,13,16,19): two distinct pathways. Prostaglandins Leukot Essent Fatty Acids 68: 181–6

Robinson CV, Rohacs T, Hansen SB (2019) Tools for Understanding Nanoscale Lipid Regulation of Ion Channels. Trends Biochem Sci 44: 795–806

Sadana R, Dessauer CW (2009) Physiological roles for G protein-regulated adenylyl cyclase isoforms: insights from knockout and overexpression studies. Neurosignals 17: 5–22

Sanabra C, Mengod G (2011) Neuroanatomical distribution and neurochemical characterization of cells expressing adenylyl cyclase isoforms in mouse and rat brain. J Chem Neuroanat 41: 43–54

Schultz JE, Natarajan J (2013) Regulated unfolding: a basic principle of intraprotein signaling in modular proteins. Trends Biochem Sci 38: 538–45

Schultz JE, Schmidt BH (1987) Treatment of rats with thyrotropin (TSH) reduces the adrenoceptor sensitivity of adenylate cyclase from cerebral cortex. Neurochem Int 10: 173–8

Seth A, Finkbeiner M, Grischin J, Schultz JE (2020) Gsalpha stimulation of mammalian adenylate cyclases regulated by their hexahelical membrane anchors. Cell Signal 68: 109538

Shin JJ, Loewen CJ (2011) Putting the pH into phosphatidic acid signaling. BMC Biol 9: 85

Sinha SC, Sprang SR (2006) Structures, mechanism, regulation and evolution of class III nucleotidyl cyclases. Rev Physiol Biochem Pharmacol 157: 105–40

Soto-Velasquez M, Hayes MP, Alpsoy A, Dykhuizen EC, Watts VJ (2018) A Novel CRISPR/Cas9-Based Cellular Model to Explore Adenylyl Cyclase and cAMP Signaling. Mol Pharmacol 94: 963–972

Sunahara RK, Taussig R (2002) Isoforms of mammalian adenylyl cyclase: multiplicities of signaling. Mol Interv 2: 168–84

Tang WJ, Gilman AG (1995) Construction of a soluble adenylyl cyclase activated by Gs alpha and forskolin. Science 268: 1769–72

Tesmer JJ, Sprang SR (1998) The structure, catalytic mechanism and regulation of adenylyl cyclase. Current opinion in structural biology 8: 713–719

Tesmer JJ, Sunahara RK, Gilman AG, Sprang SR (1997) Crystal structure of the catalytic domains of adenylyl cyclase in a complex with Gsalpha.GTPgammaS. Science 278: 1907–16

Vvedenskaya O, Rose TD, Knittelfelder O, Palladini A, Wodke JAH, Schuhmann K, Ackerman JM, Wang Y, Has C, Brosch M, Thangapandi VR, Buch S, Zullig T, Hartler J, Kofeler HC, Rocken C, Coskun U, Klipp E, von Schoenfels W, Gross J et al. (2021) Nonalcoholic fatty liver disease stratification by liver lipidomics. J Lipid Res 62:100104

Wood CAP, Zhang J, Aydin D, Xu Y, Andreone BJ, Langen UH, Dror RO, Gu C, Feng L (2021) Structure and mechanism of blood-brain-barrier lipid transporter MFSD2A. Nature 596: 444–448

Zhang G, Liu Y, Ruoho AE, Hurley JH (1997) Structure of the adenylyl cyclase catalytic core. Nature 386: 247–53

Zhu Z, Tan Z, Li Y, Luo H, Hu X, Tang M, Hescheler J, Mu Y, Zhang L (2015) Docosahexaenoic acid alters Gsalpha localization in lipid raft and potentiates adenylate cyclase. Nutrition 31: 1025–30

Ziegler M, Bassler J, Beltz S, Schultz A, Lupas AN, Schultz JE (2017) A novel signal transducer element intrinsic to class IIIa and IIIb adenylate cyclases. FEBSJ 284: 1204–1217

